# Theta Oscillations in the Human Medial Temporal Lobe during Real World Ambulatory Movement

**DOI:** 10.1101/078055

**Authors:** Zahra M. Aghajan, Peter Schuette, Tony Fields, Michelle Tran, Sameed Siddiqui, Nick Hasulak, Thomas K. Tcheng, Dawn Eliashiv, John Stern, Itzhak Fried, Nanthia Suthana

**Affiliations:** Department of Psychiatry and Biobehavioral Sciences, Semel Institute for Neuroscience and Human Behavior, University of California, Los Angeles.; Department of Neurology, University of California, Los Angeles.; Department of Neurosurgery, David Geffen School of Medicine and Semel Institute for Neuroscience and Human Behavior, University of California, Los Angeles.; NeuroPace, Inc., Mountain View, CA; Functional Neurosurgery Unit, Tel-Aviv Medical Center and Sackler School of Medicine, Tel-Aviv University, Tel-Aviv, Israel.; Department of Psychology, University of California, Los Angeles.

## Abstract

Theta oscillations play a critical role in learning and memory by coordinating the spiking activity of neuronal ensembles via mechanisms such as spike timing dependent plasticity^1–7^. This rhythm is present in rodents where it is continuously evident during movement at frequencies within 6-12Hz^8,9^. In humans, however, the presence of continuous theta rhythm has been elusive; indeed, a functionally similar theta is thought to occur at lower frequency ranges (3-7Hz) and in shorter bouts^10–12^. This lower frequency theta rhythm is observed during a variety of behaviors, including virtual navigation, but has never been tested during **real world ambulatory movement**. Here we examined the oscillatory properties of theta within the human medial temporal lobe (MTL) in freely moving human subjects chronically implanted with the clinical NeuroPace RNS^®^ responsive neurostimulator device, capable of wireless recordings of continuous intracranial deep brain electroencephalographic (iEEG) activity. MTL iEEG recordings, together with sub-millimeter position tracking, revealed the presence of high frequency theta oscillations (6-12Hz) during ambulation. The prevalence of these oscillations was increased during fast movement compared to slow movement. These theta bouts, although occurring more frequently, were not significantly different in durations during fast versus slow movements. In a rare opportunity to study one subject with congenital blindness, we found that both the prevalence and duration of theta bouts were much greater than those in sighted subjects. Our results suggest that higher frequency theta indeed exists in humans during movement providing critical support for conserved neurobiological mechanisms for spatial navigation. The precise link between this pattern and its behavioral correlates will be an exciting area for future studies given this novel methodology for simultaneous motion capture and long term chronic recordings from deep brain targets during ambulatory human behavior.

---

The crucial role of the medial temporal lobe (MTL) in declarative memory and encoding new experiences is unequivocal based on an abundant body of literature in humans and many mammalian species^13–15^. It is posited that the temporal organization of neural assemblies in this region occurs due to the ongoing rhythmic oscillatory activity in the local field potential (LFP) through modification of synaptic connections. Additionally, various frequency bands are thought to be involved in different brain functions^16,17^. In particular, the theta rhythm—a slow oscillatory activity in the 6-12Hz frequency range—has been associated with exploratory behavior^2^, as well as REM sleep^18,19^, and different features of this rhythm have been linked to memory performance on various tasks^4,20–22^.

In the rodent hippocampus, higher frequency theta oscillations (6-12Hz) are most prominent during locomotion, whereas lower frequency theta oscillations (3-7Hz) are present durin gimmobility periods^7^. Theta has also been reported in other species such as cats^23^, and in shorter oscillatory bouts in bats^24,25^, non-human primates^26,27^, and humans^10,28^. This issue is confounded by the fact that iEEG studies in primates, contrary to those performed in rodents, have been typically done using virtual navigation in stationary subjects due to restrictions imposed by recording techniques. In light of recent rodent hippocampal recordings during virtual navigation—demonstrating significant differences in theta dynamics between virtual and real world navigation, including lower frequencies and a lack of frequency-speed dependence in virtual navigation^29,30^—it is essential to directly probe the LFP during natural voluntary human movement. In this study, we therefore conducted an experiment to elucidate the properties of theta oscillations in the MTL of freely moving humans.

Subjects were four neurosurgical patients (one congenitally blind) chronically implanted with the FDA approved NeuroPace RNS^®^ (Fig. 1a) system for treatment of epilepsy. For subject demographics see Extended Data Table 1, 2. The RNS system continuously recorded MTL iEEG activity while subjects performed a task in which they were instructed to walk along linear and circular paths at slow and fast speeds. A trial consisted of the following four movements: slow movement in straight lines; slow movement in circles; fast movement in straight lines; and fast movements in circles and the order of these instructions were randomized (Fig. 1b, see Methods). This strategy was used to ensure a wide range of movement speeds. Motion tracking, using the OptiTrack system capable of recording subjects’ positional and rotational information (Fig. 1c, d, Supplementary Video 1), was recorded simultaneously with iEEG data directly from the MTL (Fig. 2a, Supplementary Video 2, 3; see Methods). The locations of electrodes were determined by co-registration of high-resolution post-operative CT images with pre-operative high-resolution magnetic resonance images (MRI) along with automated software for MTL subregion segmentation to facilitate visualization (Fig. 2b, Extended Data Table 3, see Methods). To eliminate epochs with putative epileptic activity we used a thresholding algorithm similar to methods previously described^31^ (see Methods, Extended Data Fig. 1a), which resulted in discarding 3% of the data (median, [25^th^, 75^th^]=3.04, [1.32, 4.62]%; Extended Data Fig. 1b)

**Figure 1:**
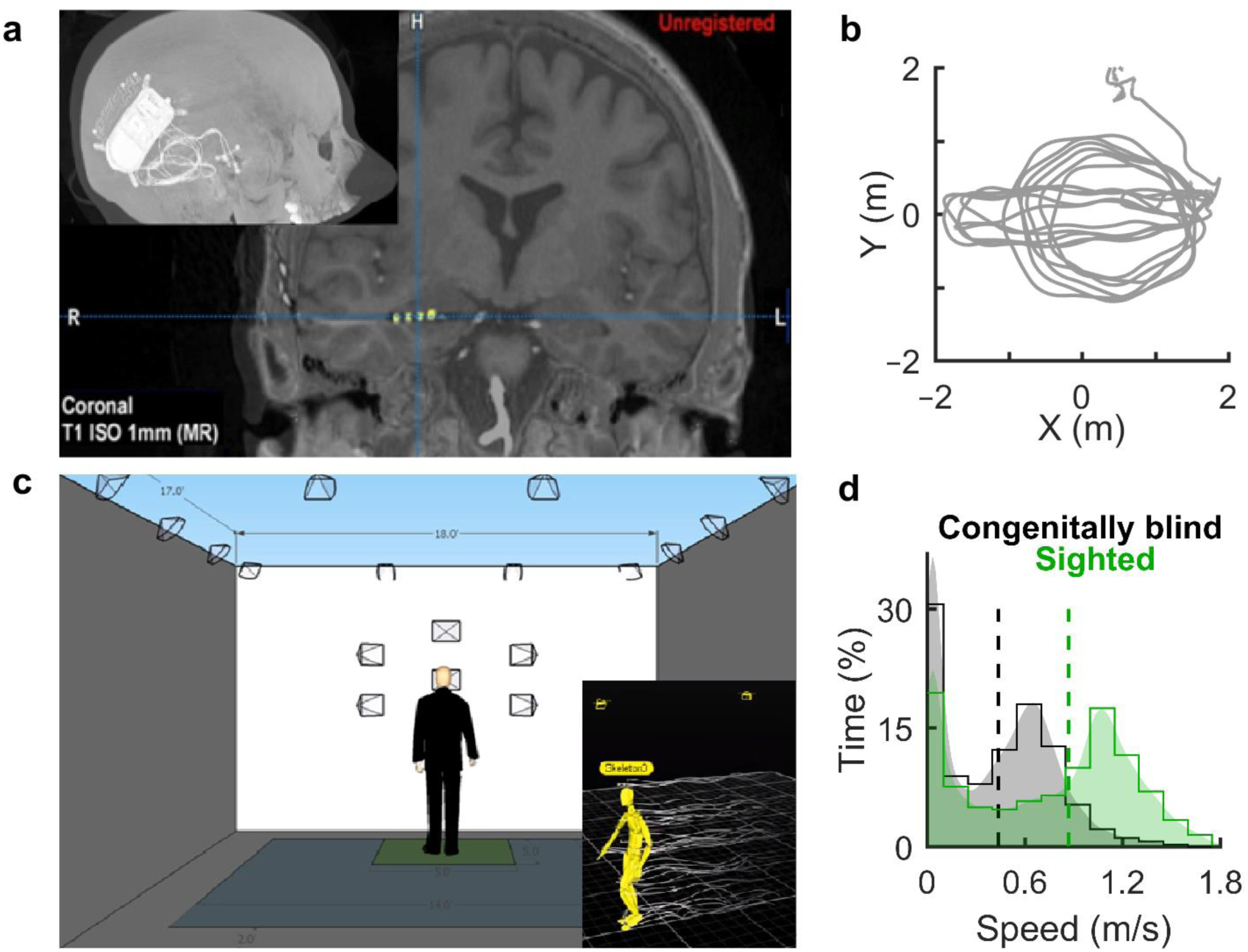
Simultaneous motion tracking and iEEG recording within the human MTL. **a,** Example post-operative CT of a subject with an implanted electrode in the right hippocampus (top left) along with a coronal view of a high-resolution MRI overlaid with co-registered electrodes (shown in yellow). **b**, Sample trajectory of a subject during one trial consisting of linear and circular movements. **c**, Schematic of the setup of cameras used for motion capture (see Methods). Inset: Real time motion tracking of an example participant. **d**, Movement speed distribution of all sighted subjects (green; median, [25^th^, 75^th^] = 0.87, [0.20, 1.14] m/s and congenitally blind subject (black; median, [25^th^, 75^th^] = 0.44, [0.05, 0.68] m/s); dashed vertical lines indicate median value; shaded areas correspond to kernel smoothing function estimates of the distributions).

**Figure 2:**
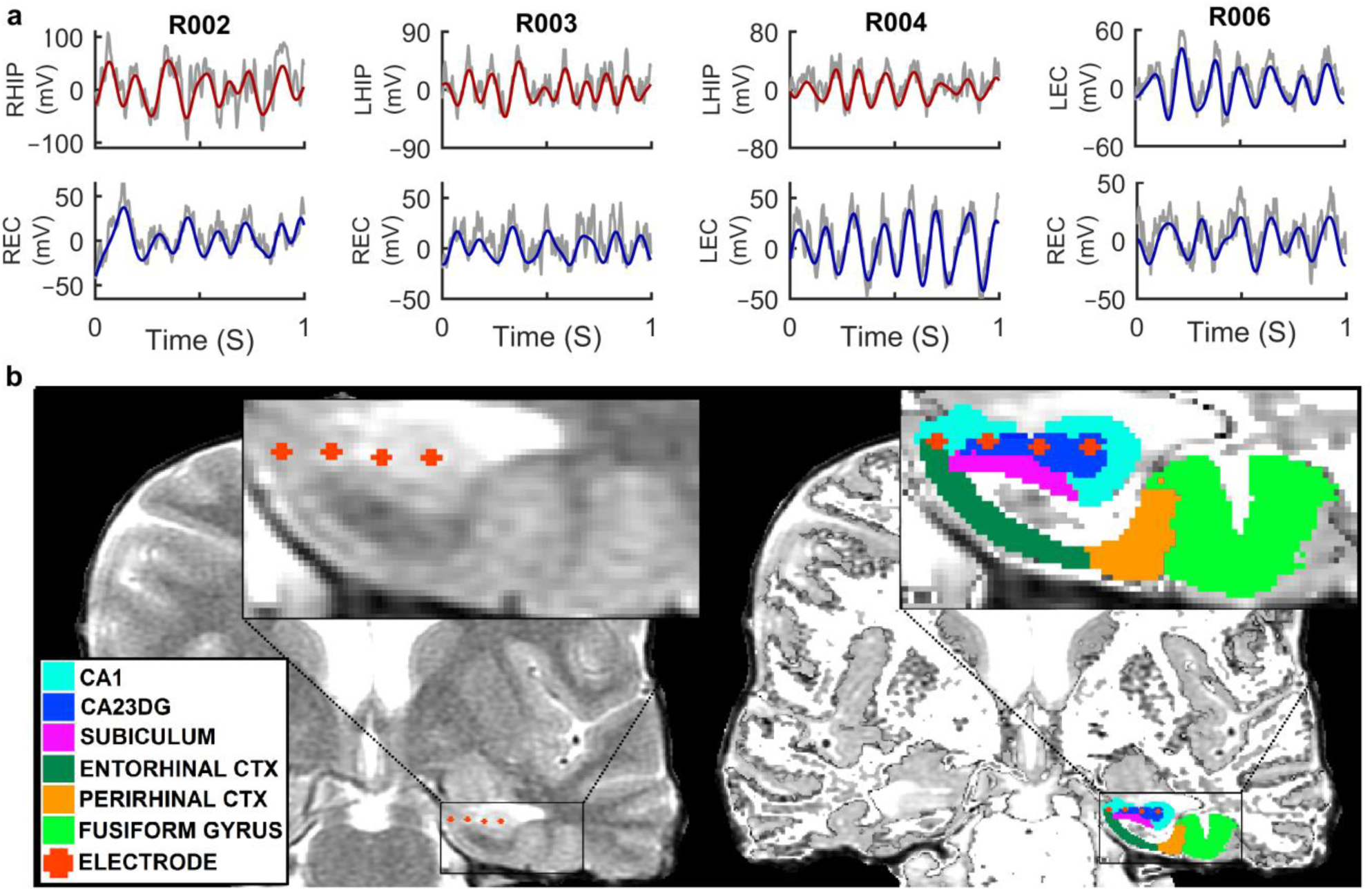
Local field potential and theta oscillations in the human MTL. **a,** Example one-second-long raw LFP traces (gray) from the MTL of all study subjects overlaid with filtered (3-12Hz) theta oscillations (hippocampal and entorhinal theta shown in red and blue respectively). **b,** Left) Sample electrode locations from a subject (R003) overlaid onto coronal pre-operative high-resolution MRI. Right) Automated MTL subregion segmentation (note that different colors correspond to different areas) demonstrating electrode locations in an example subject. White areas correspond to white matter.

Examination of raw LFP traces revealed striking theta oscillations, which were readily visible (Fig. 2a, Supplementary Video 2, 3). To further investigate this, for each subject and within each trial, data was separated into low and high speed movements using a median split on speed in that trial to obtain equal amount of data within each condition (see Methods). By utilizing the BOSC method^32^, episodes with significant oscillations (P-episodes) between 3-30 Hz (occurring for at least 3 cycles and above 95% chance level, Fig. 3a) were calculated (see Methods, Extended Data Fig. 2). Data from all trials and channels were collapsed across the frequency domain. We found that in all 4 subjects, there was a significant increase in theta oscillations, typically associated with rodent movement, during fast movements compared to slow movements (Fig. 3b). This increase in the prevalence of theta, as quantified by p-episodes, was observed between 7-9Hz (N_trials×channels_=84, p<0.05, Wilcoxon rank-sum test) for the sighted subjects and 6.5-9Hz in the congenitally blind subject (N_trials×channels_=28, p<0.05, Wilcoxon rank-sum test). Interestingly, these theta episodes were transient and present ~10% of the time for the sighted group, while this percentage was ~30% in the congenitally blind subject (Fig. 3b). Hence, we analyzed the data from the congenitally blind subject separately from the data from sighted subjects within which our results were consistent and qualitatively similar (Extended Data Fig. 3).

**Figure 3:**
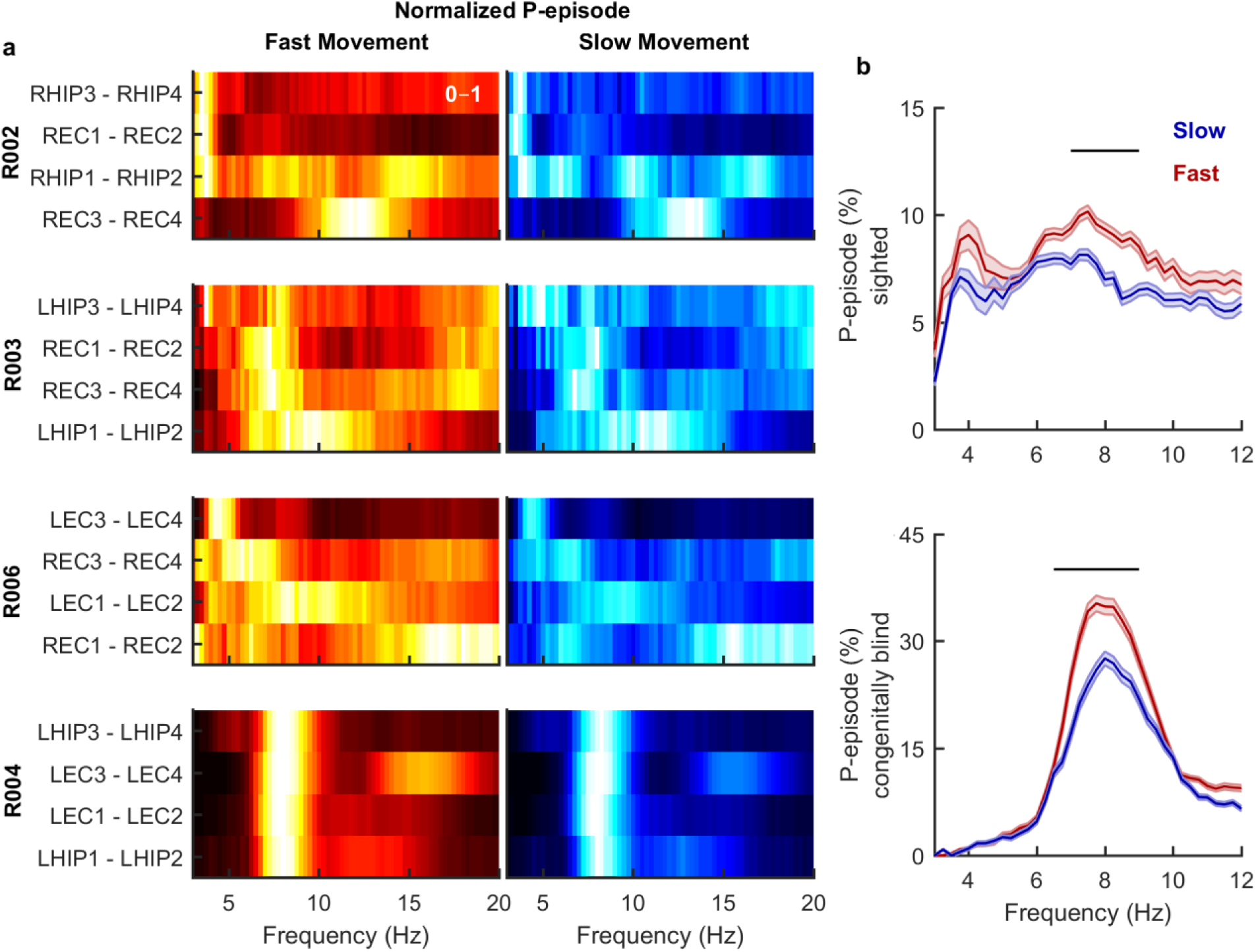
Significant increase in the prevalence of high-frequency theta oscillations during fast compared to slow movement. **a,** Colormap shows percentage of time with significant oscillations (P-episode) in the frequency range indicated on the x axis averaged across trials for each clinically labeled channel in the left and right entorhinal cortex (LEC, REC) and left and right hippocampus (LHIP and RHIP) from all subjects (electrode 1: most distal, electrode 4: most proximal). Data are normalized by the maximum value for each channel, within each condition, for visibility purposes (range: 0-1). Brighter colors indicate larger values. Throughout this figure, red shades and blue shades correspond to fast and slow movements respectively. Also, note that for each subject, channels are sorted based on the frequency with maximum prevalence in the theta band during fast movements. **Left)** Shown are normalized P-episodes during fast movement. Lighter shades indicate higher values here and throughout figures (N_trials_ for subjects R002, R003, R006 and R004 (congenitally blind) were 7, 5, 9 and 7 respectively). **Right)** Same as (**left**) but during movement at slow speeds.**) b,** Percentage of time with significant oscillations (across all trials and channels; shown are mean ± s.e.m) for low frequencies during fast movements (red) versus slow movements (blue) in 3 sighted subjects (top, N_trials×channels_=84) and 1 congenitally blind subject (bottom, N_trials×channels_=28). Black horizontal lines indicate regions with significant difference in P-episodes (p < 0.05, Wilcoxon rank-sum test.)

We then asked whether the increase in the prevalence of theta during fast versus slow movements is due to longer theta bouts or higher rates of occurrence. To address this, we computed the duration of theta bouts by allowing the number of cycles to vary in our detection analysis (Fig. 4a). Comparisons of the durations of theta bouts, as measured by the average number of theta cycles (weighted by P-episode in each frequency bin), in fast and slow movements showed no significant difference in bout durations between the two conditions (p>0.05 at any frequency; Wilcoxon rank-sum test) (Fig. 4b). This result suggests that more prevalent theta during fast movement potentially arises as a result of short theta bouts of similar lengths occurring more frequently during fast movements. We also observed that the average number of detected theta cycles was higher in the congenitally blind subject (~4) compared to sighted subjects (~3) at the peak frequency (Fig. 4b).

**Figure 4:**
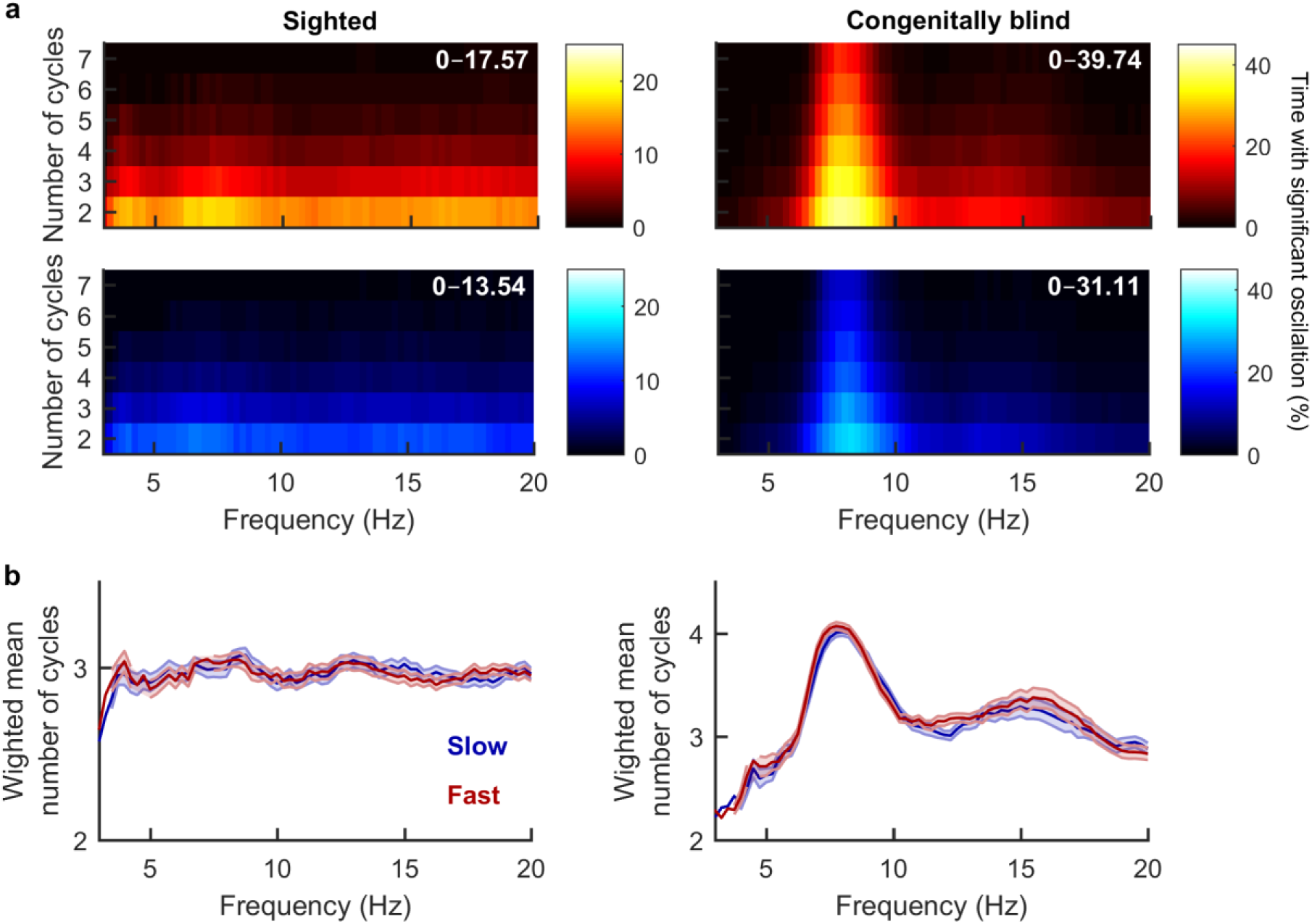
Duration of theta bouts is similar during fast and slow movements. **a,** Colormaps indicate percentage of time with significant oscillations (P-episodes) in each frequency (averaged across trials, channels and subjects) assuming varying number of minimum cycles for detection (y-axis). Red and blue color schemes correspond to fast and slow movements respectively, here and throughout the figure. Number at the top right corner indicates range. **b,** In each frequency bin, the number of detected theta cycles was weighted by the corresponding P-episode normalized by the overall P-episode for individual trials and channels to obtain weighted mean number of cycles with significant oscillations at each frequency. Shown are the mean ± s.e.m of the weighted number of theta cycles across all trials and channels. There was no significant difference between the duration of the theta bouts between fast (red) and slow (blue) movements. However, note that in the congenitally blind subject (right), the duration of theta bouts (~4 cycles at peak frequency) were longer than those in the sighted subjects (left, ~3 cycles).

Classification of behavior into fast and slow movement speeds was further tested using a neural networks machine-learning model (Extended Data Fig. 4a, see Methods). Here, power spectra—computed using BOSC method—were used as the input to our model in order to predict movement speed. Receiver operating characteristic (ROC) plots and the area under these curves (AUC) showed that the performance of our model was significantly better than chance in classifying fast from slow movement speeds (Extended Data Fig. 4b).

We present a novel innovative methodology for combining simultaneous motion capture technology and continuous iEEG recordings performed in the same subjects in a unique setting enabling access to the MTL in awake conscious freely moving humans. To our knowledge, this study demonstrates the first quantification of theta oscillations during human ambulatory movement. Our results show high frequency (~8Hz) theta oscillations occur in short bouts and significantly differentiate fast from slow movement speeds in that they are present more frequently during the former.

Previous investigations of theta oscillations in primates have resulted in conflicting evidence regarding the presence and functionality of the theta rhythm compared to those in rodents^11,12^,^27,33^. These dissimilarities are thought to be in part attributed to the differences in the primary source of sensory information in these species—visual inputs in primates versus olfactory and somatosensory inputs in rodents. Nonetheless, the lack of feasibility in recording iEEG in ambulatory primates (in contrast to rodents), has not previously allowed for unequivocal evaluation of theta oscillatory properties under similar conditions in these species. Our findings demonstrate that in humans short, intermittent, theta bouts happen at a higher rate during fast movements which could potentially be explained by a higher rate of saccadic eye movements, as the dominant source of sensory inputs, at faster speeds^27^. Whether these theta bouts are elicited and phase-reset by eye saccades warrants further investigation.

Curiously, in our present results, significant theta oscillations occur more often and more continuously in a single subject who is congenitally blind. This subject used a Hoover cane to traverse the environment, thereby possibly using somatosensory inputs to a greater extent compared to other normally sighted subjects, supporting the idea that the modality of sensory processing may be a critical factor relating theta oscillations to movement.

It is possible that there are subtle oscillatory related differences across MTL regions and along the anterior-posterior axis, which are not captured within our study. For example, in rodents, theta oscillations are diminished in power and spatial selectivity is reduced in the ventral compared to dorsal hippocampus^34^. In humans, the dorsoventral distinction is thought to map onto the posterior-anterior hippocampal axis^35^. Therefore, future larger sample studies will be necessary to characterize movement-related theta changes across MTL regions and along the anterior-posterior axis in humans.

Recently, it has been argued that the human analogue of rodent theta oscillations exists in the lower (<4Hz) frequency range^10^. However, to our knowledge, theta oscillations within the human MTL have never been investigated during real world ambulatory movement due to limitations of wired intracranial recording electrode technology. Using wireless, chronically implanted electrodes, our results suggest there does in fact exist a higher frequency (~8Hz) theta oscillation in the human MTL associated with physical movement. While the high frequency theta presented in the current study varies significantly between different speeds of movement, its exact behavioral correlates—and how it relates to memory performance—remains to be examined and tested in future experiments with memory demands. Furthermore, although lower frequency theta in humans has been observed in stationary virtual navigation and memory tasks^36–39^, future studies are needed to determine low and high frequency theta dynamics during ambulatory movement compared to those in stationary behavioral tasks. Moreover, these two patterns of theta oscillations in rodents, namely type-1 (high frequency) and −2 (low frequency), are thought to be functionally distinct with the former connected with locomotion and the latter involved in memory and learning^8,40,41^.

Overall, the current study provides important insight into human MTL theta band oscillatory dynamics during ambulatory behavior while bridging findings across species. We present a novel paradigm for the leveraging of FDA technology combined with state-of-the-art motion tracking that allows for future investigation of neural oscillatory dynamics during real world behaviors in humans.

## Methods

### Data acquisition

The FDA approved Neuropace RNS^®^ System (Fig. 1a) is designed to detect abnormal electrical activity in the brain and respond in a closed loop manner by delivering imperceptible levels of electrical stimulation to normalize brain activity before an individual experiences seizures. For the present study, a neurologist was present to configure stimulation to OFF and ON before and after the study and monitor for seizure activity. Each of the 4 subjects in the study had two implanted depth electrode leads 1.27 mm in diameter each with 4 platinum-iridium electrode contacts, each with a surface area of 7.9 mm^2^, 1.5 mm long with an electrode spacing of either 3.5 or 10 mm (Extended Data Table 3). During the study, the RNS^®^ Neurostimulator continuously monitored iEEG activity on four bipolar channels at 250 Hz with an analogue filter equivalent to a 1^st^ order Butterworth with 3db attenuation at the cutoff frequencies (4-90Hz). The neurostimulator communicates wirelessly with a Programmer and Remote Monitor using secure protocols. The Programmer is used to (1) retrieve data, including stored iEEG data, from the neurostimulator, (2) configure detection and stimulation, and (3) monitor iEEG activity and test stimulation settings in real-time. A programmable electromagnet was used to trigger iEEG storage in the RNS Neurostimulator.

### Electromagnet

A programmable electromagnet was designed to be placed on the subject’s head over the RNS Neurostimulator and trigger iEEG recordings by generating magnetic pulses. The magnetic pulses could be triggered either manually or automatically with programmable intervals of 30, 60, 90, 150, 180 and 240 seconds. Power was provided by rechargeable NICAD batteries. The electromagnet had a visible LED as well as an infrared LED, which were used both as visible indicators and event markers for synchronization of iEEG recordings with recorded video. The device accepts three commands: a hard reset command which allows for either alteration of the timing or entering into the test mode; a clear command which is a soft reset in that any command other than reset is halted and all timing counters are cleared; and a start or stop command which starts the timing for the marker function.

### Motion Tracking

Motion tracking was done using the Optitrack system (Natural Point, Inc.) with 8 ceiling mounted infrared high-resolution cameras that allow for sub-millimeter motion tracking. Removable reflective markers were used in conjunction with specialized software, Motive and Camera SDK, which allows for kinematic labeling. For the head, a rigid body object was constructed, from the reflective markers, for which center of mass (i.e. position) was tracked in addition to a quaternion describing rotational information that is translated to Euler angles to obtain yaw, pitch and roll with respect to the experimental room. This information was then used to obtain subjects’ movement trajectory and speed (Fig. 1b, c, d). Further, another set of 4 ceiling mounted high-resolution Optitrack cameras were dedicated to record videos in the visible spectrum, which were then used for synchronization purposes (see below).

### Synchronization of iEEG data and motion capture data

When the electromagnet was activated it triggered the storage of iEEG data by the RNS Neurostimulator in preconfigured short durations (60 s), generated a “magnet marker” event in the iEEG data, as well as turned on a visible LED light, which was captured by the video cameras. The “magnet marker” events in the iEEG data were then aligned with the onset of the visible LED light events in our motion tracking system. This allowed us to synchronize the two data streams and combine the stored pieces of iEEG data into a long continuous recording.

### Subjects

Subjects (Extended Data Table 1) were 4 patients (three sighted and one congenitally blind) with pharmacoresistant epilepsy who are implanted with the FDA approved NeuroPace RNS device for treatment of epilepsy. The congenitally blind subject exhibited lifelong visual impairment with marginal light perception due to retinopathy of prematurity. Electrode placements were determined solely based on clinical criteria. All subjects volunteered for the study by providing informed consent according to a protocol approved by the UCLA Medical Institutional Review Board (IRB). Neuropsychological scores for each individual were determined using methods previously described^42^(Extended Data Table 2).

### Electrode Localization

Electrode localization was done using methods similar to those reported by Suthana *et al.*^42^. A high-resolution post-operative CT image was co-registered to a pre-operative whole brain and high-resolution MRIs for each subject (Fig. 2b). We used the FSL FLIRT (FMRIB’s Linear Registration Tool^43^) together with BrainLab stereotactic and localization software^44^. MTL regions (entorhinal, perirhinal, parahippocampal, hippocampal subfields CA23DG [CA2, 3, dentate gyrus], CA1, and subiculum) are anatomically determined by boundaries that are demarcated based on atlases correlating MRI visible landmarks with underlying cellular histology. To avoid human bias and perform hippocampal segmentation programmatically we used ASHS software^45^.

### Behavioral task

Participants performed a task in which they were directed to walk slow or fast following linear and circular paths in a 400 square foot room based on an auditory command. Each trial consisted of four different conditions: walking in a straight line or circle at slow or fast speeds and the order in which these conditions were presented was randomized. This strategy yielded a relatively large range of movement speeds (Fig. 1d). Furthermore, for each subject, the speed profile was similar across different trials (data not shown).

### Data Analysis

All analyses were done offline using custom codes in MATLAB.

A. Elimination of epileptic activity from LFP: Data from putative epileptic epochs were discarded according to a method similar to procedures described recently^31^ using functions found in MATLAB Signal Processing Toolbox. In brief, epileptic discharges were identified when either of the following conditions were satisfied: a) the envelope of the unfiltered signal was 5 s.d. above the baseline; the envelope of the filtered signal (band-pass filtered in the 25-80Hz range followed by signal rectification) was 6 s.d. above the baseline (Extended Data Fig. 1). We excluded ~3% of the data from further analysis because of the presence of epileptic activity and this percentage was not significantly different between slow and fast movements (p = 0.6, Wilcoxon rank-sum test; Extended Data Fig. 1b).
B. Detection of significant oscillations within different frequency ranges: BOSC algorithm was used^32^ and episodes with significant oscillations between 3-12Hz (using 6^th^ order wavelets and for bouts occurring for at least 3 cycles and above 95% chance level) were detected. Additionally, to examine the duration of theta bouts, the lower bound on the number of cycles for detection was allowed to vary.
C. Control analysis for computing power spectra and oscillation detection: To evaluate the effect of the analog bandpass filter (1^st^ order Butterworth 4-90Hz, 3db attenuation at cutoff frequencies) on the RNS System on our results, we mathematically implemented this filter on data collected from a previous study^42^ in patients with implanted depth electrodes. We first computed the power spectrum, using BOSC method, for randomly selected (N=100) 5-seconds-long LFPs during trials (P_1_). Power spectra were then computed for these LFPs filtered using the abovementioned filter settings (P_2_). A similarity index was computed between the two matrices (P_1_ and P_2_) using cosine distance defined as follows:

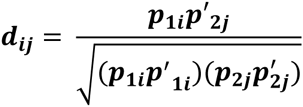 A similarity index was then calculated while allowing for the frequency lower bound to vary (Extended Data Fig. 2a). The same procedure was done on the similarity index calculated between the matrices consisting of data points with significant oscillations (Extended Data Fig. 2b).
D. Extraction of theta from LFP: Raw signal was bandpass filtered (between 3-12Hz) using an acausal 4^th^ order Butterworth filter (Fig. 2a). The phase and amplitude of theta was then computed from the filtered signal using Hilbert transform.
E. Statistics: Two-sided nonparametric Wilcoxon rank-sum test was utilized to assess the significant differences between linear variables. For each histogram, a kernel smoothing function was also computed for the probability density estimates. Values are expressed as median, [25^th^, 75^th^] or mean ± s.e.m when applicable.
F. Classification of movement speed using machine learning: To classify behavior into fast and slow movement speeds (here referring to the top and bottom 30% of the running speeds respectively), we used a feed forward neural network with an input layer, a hidden layer consisting of 100 neurons and an output layer with a binary classifier (the corresponding speed categories). A tan-sigmoid function and a scaled conjugate gradient algorithm were used as a transfer function and back propagation algorithm respectively (Extended Data Fig. 4a). For each subject, the BOSC method was utilized to compute a power spectrum of the LFP for each time point. Data from all channels were concatenated (leaving a contiguous 15% of the total data out for the testing set to ensure independence). For the remaining data, 85% was used for training while the other 15% was used for cross validation. Receiver operating characteristic (ROC) plot and the area under that curve (AUC) were used to evaluate the performance of our model (Extended Data Fig. 4b).

## Acknowledgements

This work was supported by UCLA startup funds (PI: Suthana) and the DARPA Restoring Active Memory program (Agreement Number: N66001-14-2-4029). We thank Eric Behnke for technical assistance; Brooke Salaz for general assistance, Fabien Scalzo for useful discussions; and the subjects for their participation in this study.

## Author Contributions

Z.M.A. and N.S., designed research; Z.M.A, P.S., M.T., S.S. and N.S. performed analyses; Z.M.A, P.S., and N.S. collected data and performed research; Z.M.A., T.F, N.H., T.T., and N.S. designed and implemented technical tools; D.E., J.S., and I.F. provided clinical and neurosurgical procedures; and Z.M.A, I.F., and N.S. wrote the paper. All the authors commented on the manuscript.

## Author information

Reprints and permission information is available at www.nature.com/reprints. T.T and N.H. are employees of NeuroPace, Inc. The authors have no other relevant affiliations or financial involvement with any organization or entity with a financial interest in or financial conflict with the subject matter or materials discussed in the manuscript apart from those disclosed. Correspondence should be addressed to N.S.

---

**Extended Data Figure 1:**
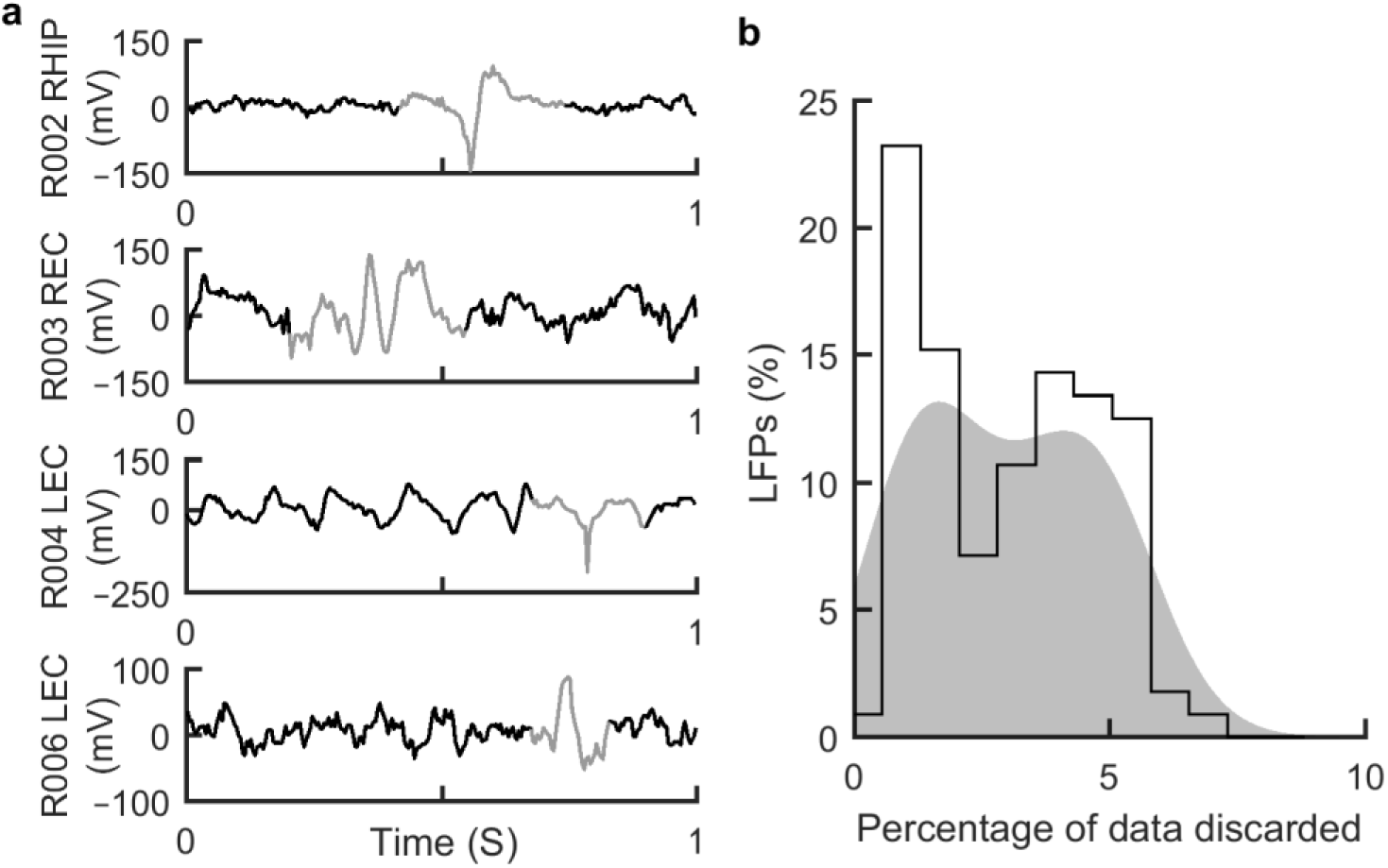
Elimination of epileptic discharges. **a)** Shown are example one-second-long LFP traces from 4 different subjects (in black) and epileptic discharges detected (in grey) using thresholding methods (see Methods). **b)** Putative epileptic epochs were detected on LFPs from all channels and within all trials, and data from these periods were discarded from analysis (percentage of epileptic epochs = 3.04, [1.32, 4.62]%, N_trials×channels_ = 112). The percentage of putative epileptic data was similar during slow movements (3.08, [1.37, 4.41]%) and fast movements (3.01, [1.46, 5.01]%) and not significantly different from one another (p = 0.6, Wilcoxon rank-sum test). Numbers are reported as median, [25th, 75th]. Shaded area corresponds to kernel smoothing function estimate of the distribution.

**Extended Data Figure 2:**
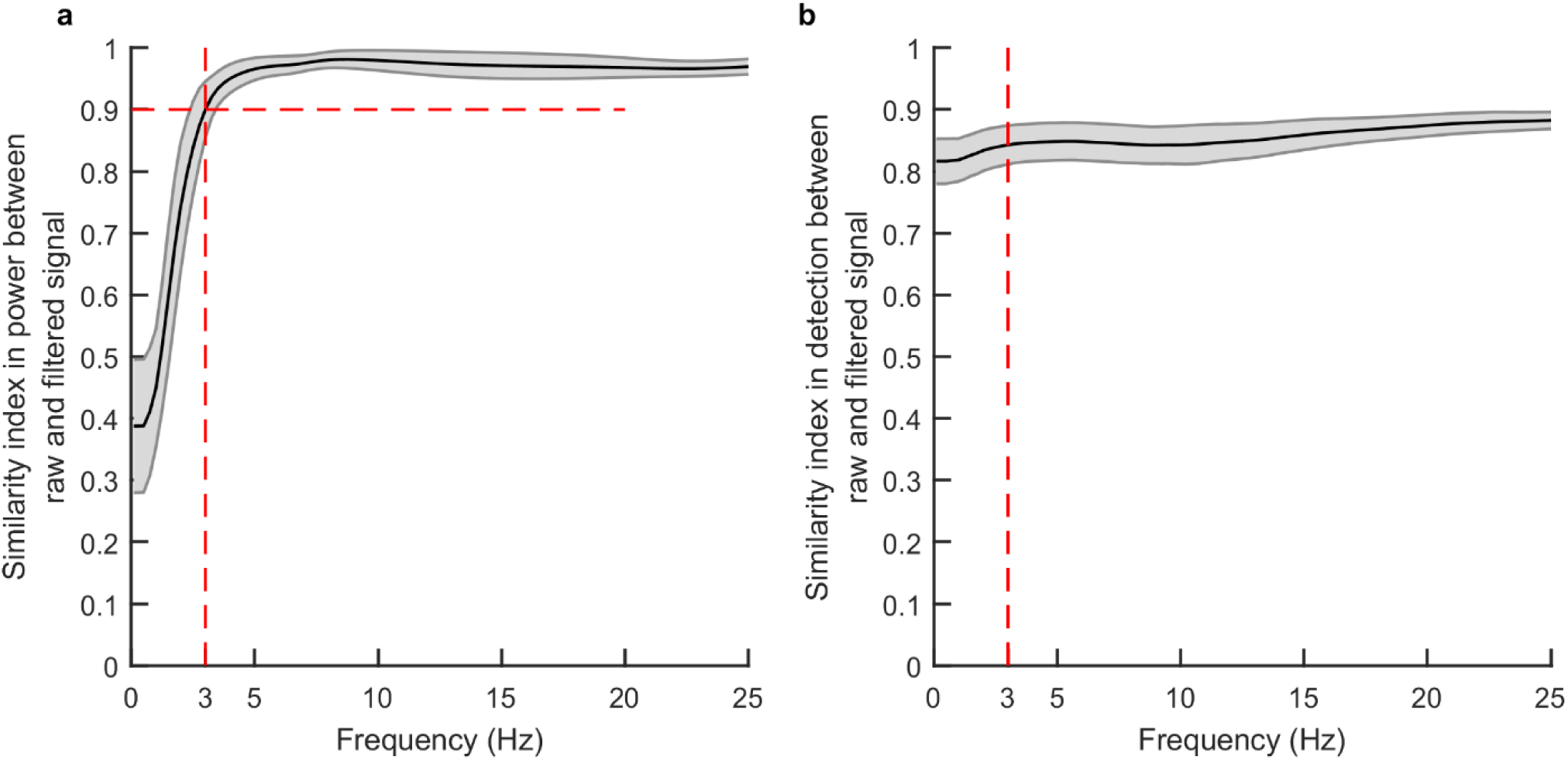
Control analysis for comparisons of raw wide-band signal and filtered signal in oscillation detection methods. We used data collected from a previous study42 in patients implanted with intracranial depth electrode for seizure monitoring. We mathematically implemented a filter, analogous to that existing on the RNS system, on this wide-band data to investigate the effect of filtering in our oscillation detection algorithms. **a,** Similarity index between power spectra computed using BOSC methods from the raw wide-band signal and bandpass filtered signal after the frequencies shown on x-axis (mean ± s.e.m, N = 100). Dashed red line indicates the frequency above which the similarity index is higher than 0.9 (F = 3Hz). **b,** Similarity index between the matrices consisting of significant oscillations detected in the raw and filtered signal. After F=3Hz, this index is above 0.84 and consequently we used this value (3 Hz) as the minimum value for interpreting our results.

**Extended Data Figure 3:**
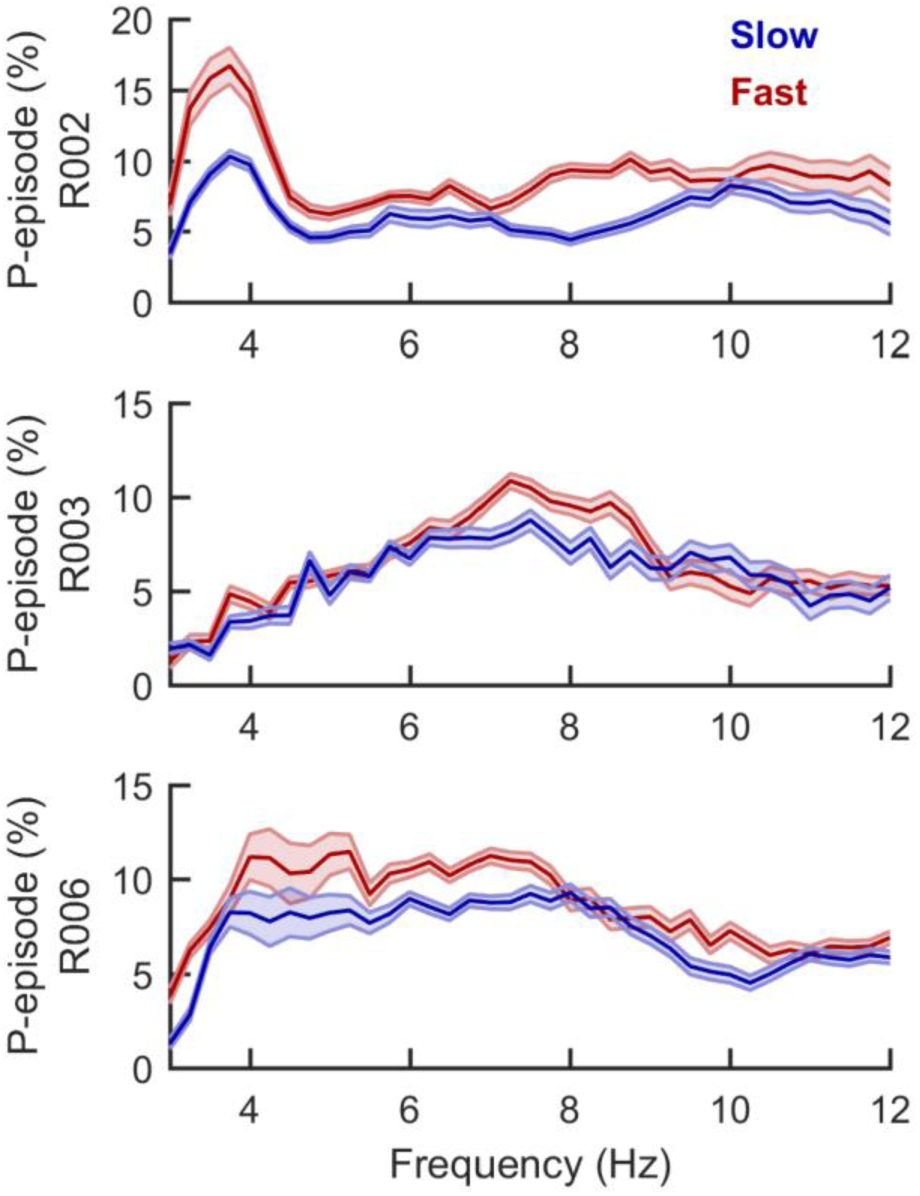
Increased prevalence of theta oscillations during fast versus slow movements within individual sighted subjects. Analysis of p-episodes within each sighted subject (individual rows) exhibited qualitatively similar results to the overall results from all subjects shown in Fig. 3b. Namely, theta was more prevalent during fast (red) compared to slow (blue) movements. Shown are the mean ± s.e.m).

**Extended Data Figure 4:**
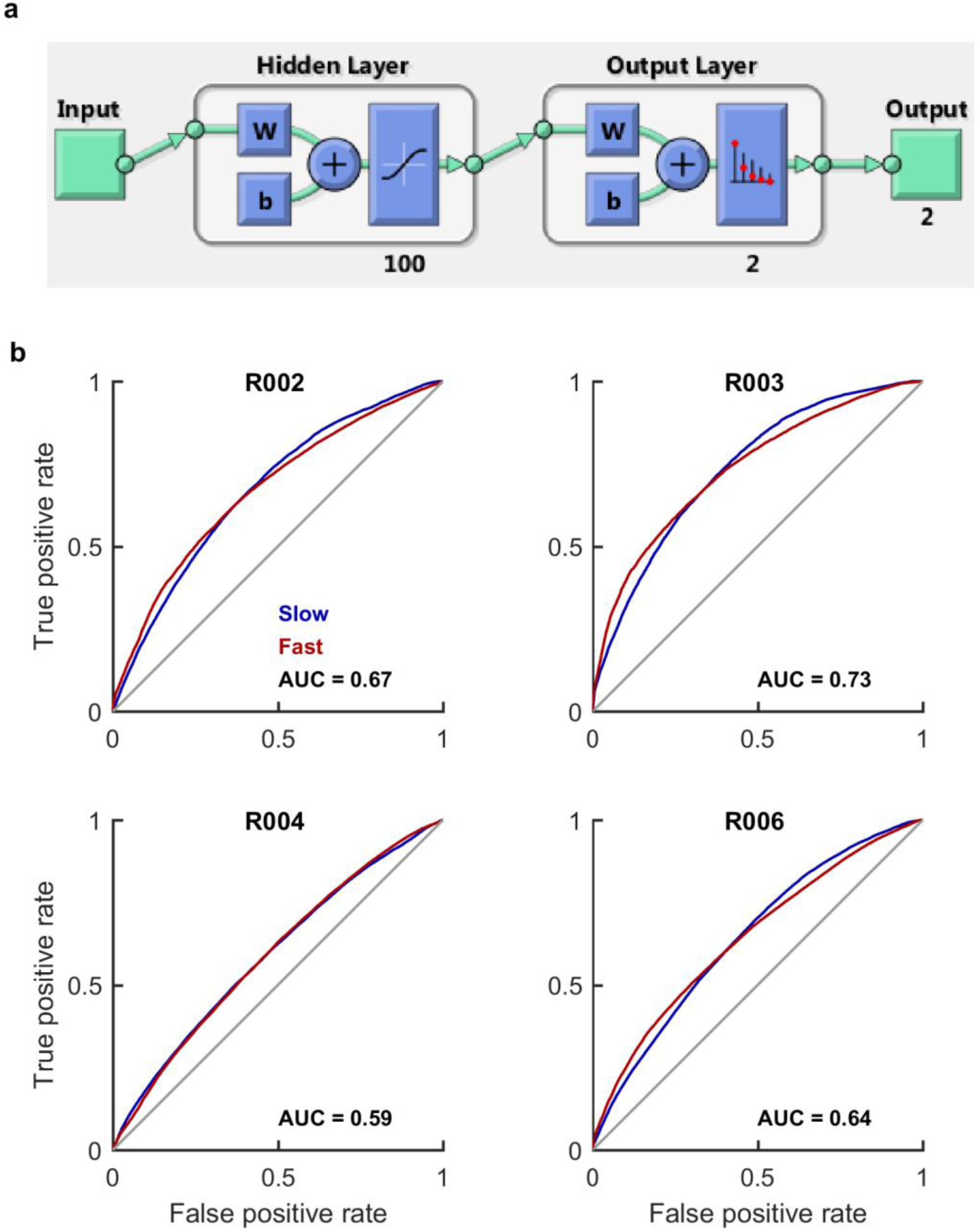
Neural networks as a tool to decode movement speed. **a)** The structure of the network utilized for speed classification consisting of 100 neurons in the hidden layer and two output classes corresponding to the slow and fast movement speeds. Input data was the power spectra in 3-30Hz frequency band computed using BOSC method (see Methods) and concatenated across all 4 channels. Classification was done separately for each subject. **b)** ROC plots from all subjects showed that our model could successfully predict the two classes (slow speeds in red, fast speeds in blue) as indicated by the distance between the lines from the two classes and chance level (grey diagonal line). The area under the ROC curve (AUC), a measure commonly used to describe the performance of a classification model, for each subject is reported at the bottom right. Note that in all case, AUC values are above 0.5 (the area under diagonal line or the chance level).

**Extended Data Table 1:** Subject demographics.

| **Subject** | **Age** | **Gender** | **Handedness** |
| --- | --- | --- | --- |
| R002 | 34 | F | R |
| R003 | 40 | F | L |
| R004* | 63 | M | R |
| R005 | 40 | F | R |
| R006 | 45 | M | L |

Demographics of the study subjects including age, gender and handedness. Data from Subject R005 was excluded from analysis due to epileptic activity.
\* Subject R004 is congenitally blind.

**Extended Data Table 2:** Clinical Characteristics of the Study Subjects

| Subject No. | Verbal IQ | Digit Span | Verbal Memory |  | Visual Memory | Executive Function |
| --- | --- | --- | --- | --- | --- | --- |
|  |  |  | WMS | CVLT<br>*Percentile* |  |  |
| R002 | 113 | 10 | 66 | 79 | 34.5 | 91 |
| R003 | 70 | 37 | <1 | <1 | 2 | 25 |
| R004 | 108 | 14 | 73 | 84 | - | 84 |
| R006 | NA | 16 | <1 | <1 | <1 | 66 |

Each subject performed neuropsychological tests for clinical characteristics evaluation using methods described earlier<sup>42</sup>. In brief, Wechsler Adult Intelligence Scale was used to calculate verbal and digit span. Verbal memory was assessed by the logical memory portion of the Wechsler Memory Scale (WMS) and the long-delay free-recall part of the California Verbal Learning Test (CVLT). Visual memory and executive function were tested using Rey-Osterrieth Complex Figure test and the Trail Making Test respectively. All measures are reported as percentiles with the exception of Verbal IQ reported as standardized scores.

**Extended Data Table 3:** Electrode Localizations.

| Subject | Electrode Labels | Electrode Spacing | Localization |
| --- | --- | --- | --- |
| R002 | REC1 | 10 mm | Entorhinal Cortex |
|  | REC2 |  | Perirhinal Cortex |
|  | REC3 |  | Fusiform Gyrus |
|  | REC4 |  | Inferior Temporal |
|  | RHIP1 | 3.5 mm | CA1 |
|  | RHIP2 |  | CA1 |
|  | RHIP3 |  | CA1 |
|  | RHIP4 |  | CA1 |
| R003 | REC1 | 3.5 mm | Entorhinal Cortex |
|  | REC2 |  | Perirhinal Cortex |
|  | REC3 |  | Perirhinal Cortex |
|  | REC4 |  | Fusiform Gyrus |
|  | LHIP1 | 3.5 mm | CA1 |
|  | LHIP2 |  | CA23DG |
|  | LHIP3 |  | CA23DG |
|  | LHIP4 |  | CA23DG |
| R004 | LEC1 | 10 mm | Entorhinal Cortex |
|  | LEC2 |  | Perirhinal Cortex |
|  | LEC3 |  | Inferior Temporal |
|  | LEC4 |  | Inferior Temporal |
|  | LHIP1 | 3.5 mm | CA1 |
|  | LHIP2 |  | CA1 |
|  | LHIP3 |  | CA1 |
|  | LHIP4 |  | CA1 |
| R006 | LEC1 | 3.5 mm | Entorhinal Cortex |
|  | LEC2 |  | Perirhinal Cortex |
|  | LEC3 |  | Perirhinal Cortex |
|  | LEC4 |  | Fusiform Gyrus |
|  | REC1 | 3.5 mm | Subiculum |
|  | REC2 |  | Subiculum |
|  | REC3 |  | CA1 |
|  | REC4 |  | Fusiform Gyrus |

## Supplementary Information

**Supplementary video 1: Real time motion tracking of an example subject**

**Supplementary video 2: Simultaneous movement and MTL LFP recordings (example sighted subject)**

**Top)** Movement trajectory of a subject during motion along a linear path (shown in black) and the subject’s head direction indicated by red arrow. **Bottom)** Continuous LFP recording simultaneously during the motion shown above.

**Supplementary video 3: Simultaneous movement and MTL LFP recordings (congenitally blind subject)**

**Top)** Movement trajectory of a subject during motion along a linear path (shown in black) and the subject’s head direction indicated by red arrow. **Bottom)** Continuous LFP recording simultaneously during the motion shown above.

